# The Genomic Formation of South and Central Asia

**DOI:** 10.1101/292581

**Authors:** Vagheesh M. Narasimhan, Nick Patterson, Priya Moorjani, Iosif Lazaridis, Mark Lipson, Swapan Mallick, Nadin Rohland, Rebecca Bernardos, Alexander M. Kim, Nathan Nakatsuka, Iñigo Olalde, Alfredo Coppa, James Mallory, Vyacheslav Moiseyev, Janet Monge, Luca M. Olivieri, Nicole Adamski, Nasreen Broomandkhoshbacht, Francesca Candilio, Olivia Cheronet, Brendan J. Culleton, Matthew Ferry, Daniel Fernandes, Beatriz Gamarra, Daniel Gaudio, Mateja Hajdinjak, Éadaoin Harney, Thomas K. Harper, Denise Keating, Ann Marie Lawson, Megan Michel, Mario Novak, Jonas Oppenheimer, Niraj Rai, Kendra Sirak, Viviane Slon, Kristin Stewardson, Zhao Zhang, Gaziz Akhatov, Anatoly N. Bagashev, Bauryzhan Baitanayev, Gian Luca Bonora, Tatiana Chikisheva, Anatoly Derevianko, Enshin Dmitry, Katerina Douka, Nadezhda Dubova, Andrey Epimakhov, Suzanne Freilich, Dorian Fuller, Alexander Goryachev, Andrey Gromov, Bryan Hanks, Margaret Judd, Erlan Kazizov, Aleksander Khokhlov, Egor Kitov, Elena Kupriyanova, Pavel Kuznetsov, Donata Luiselli, Farhod Maksudov, Christopher Meiklejohn, Deborah Merrett, Roberto Micheli, Oleg Mochalov, Zahir Muhammed, Samariddin Mustafokulov, Ayushi Nayak, Rykun M. Petrovna, Davide Pettener, Richard Potts, Dmitry Razhev, Stefania Sarno, Kulyan Sikhymbaeva, Sergey M. Slepchenko, Nadezhda Stepanova, Svetlana Svyatko, Sergey Vasilyev, Massimo Vidale, Dmitriy Voyakin, Antonina Yermolayeva, Alisa Zubova, Vasant S. Shinde, Carles Lalueza-Fox, Matthias Meyer, David Anthony, Nicole Boivin, Kumarasamy Thangaraj, Douglas J. Kennett, Michael Frachetti, Ron Pinhasi, David Reich

## Abstract

The genetic formation of Central and South Asian populations has been unclear because of an absence of ancient DNA. To address this gap, we generated genome-wide data from 362 ancient individuals, including the first from eastern Iran, Turan (Uzbekistan, Turkmenistan, and Tajikistan), Bronze Age Kazakhstan, and South Asia. Our data reveal a complex set of genetic sources that ultimately combined to form the ancestry of South Asians today. We document a southward spread of genetic ancestry from the Eurasian Steppe, correlating with the archaeologically known expansion of pastoralist sites from the Steppe to Turan in the Middle Bronze Age (2300-1500 BCE). These Steppe communities mixed genetically with peoples of the Bactria Margiana Archaeological Complex (BMAC) whom they encountered in Turan (primarily descendants of earlier agriculturalists of Iran), but there is no evidence that the main BMAC population contributed genetically to later South Asians. Instead, Steppe communities integrated farther south throughout the 2nd millennium BCE, and we show that they mixed with a more southern population that we document at multiple sites as outlier individuals exhibiting a distinctive mixture of ancestry related to Iranian agriculturalists and South Asian hunter-gathers. We call this group *Indus Periphery* because they were found at sites in cultural contact with the Indus Valley Civilization (IVC) and along its northern fringe, and also because they were genetically similar to post-IVC groups in the Swat Valley of Pakistan. By co-analyzing ancient DNA and genomic data from diverse present-day South Asians, we show that *Indus Periphery*-related people are the single most important source of ancestry in South Asia—consistent with the idea that the *Indus Periphery* individuals are providing us with the first direct look at the ancestry of peoples of the IVC—and we develop a model for the formation of present-day South Asians in terms of the temporally and geographically proximate sources of *Indus Periphery-*related, Steppe, and local South Asian hunter-gatherer-related ancestry. Our results show how ancestry from the Steppe genetically linked Europe and South Asia in the Bronze Age, and identifies the populations that almost certainly were responsible for spreading Indo-European languages across much of Eurasia.

**One Sentence Summary:** Genome wide ancient DNA from 357 individuals from Central and South Asia sheds new light on the spread of Indo-European languages and parallels between the genetic history of two sub-continents, Europe and South Asia.

## Main text

### Ancient DNA Data and Analysis Strategy

We generated whole-genome ancient DNA data from 362 previously unreported ancient individuals and higher quality data from 17 previously reported individuals. Almost all derive from three broad regions: 132 from Iran and the southern part of Central Asia (present-day Turkmenistan, Uzbekistan, and Tajikistan, which we call Turan; “Iran/Turan”), 165 from the western and central Steppe and northern forest zone encompassing present day Kazakhstan and Russia (“Forest Zone/Steppe”), and 65 from northern Pakistan (“South Asia”). Our dataset includes the first published ancient DNA data from 1) Chalcolithic and Bronze Age eastern Iran and Turan (5600-1200 BCE from 12 sites); 2) early ceramic-using hunter-gatherers from the western Siberian forest zone (6200-4000 BCE from 2 sites); 3) Chalcolithic and Bronze Age pastoralists from the Steppe east of the Ural mountains, including the first ancient data from Bronze Age Kazakhstan (4700-1000 BCE from 20 sites); and 4) the first ever ancient DNA from South Asia from Iron Age and historical settlements in the Swat Valley of Pakistan (1200 BCE – 1 CE from 7 sites) (**Fig. 1, Supplementary Materials, Data S1**). To generate these data, we prepared samples in dedicated clean rooms, extracted DNA (*1, 2*), constructed libraries for Illumina sequencing (*3, 4*), and screened them using previously described procedures (*5–7*). We enriched the libraries for DNA overlapping around 1.24 million single nucleotide polymorphisms (SNPs), sequenced the products on Illumina instruments, and performed quality control (**Data S1**) (*5, 6, 8*). We also report 186 new direct radiocarbon dates on human bone (**Data S2**). After grouping individuals based on archaeological and chronological information and merging with previously reported data, our dataset included 612 ancient individuals that we then co-analyzed with genome-wide data from present-day individuals genotyped at around 600,000 SNPs, 1,789 of which were from 246 ethnographically-distinct groups in South Asia (**Data S3; Supplementary Materials**) (*9–11*). We restricted analyses to ancient samples covered by at least 15,000 SNPs. We use *Italic* font to refer to genetic groupings and normal font to indicate archaeological cultures or sites.

**Fig. 1.**
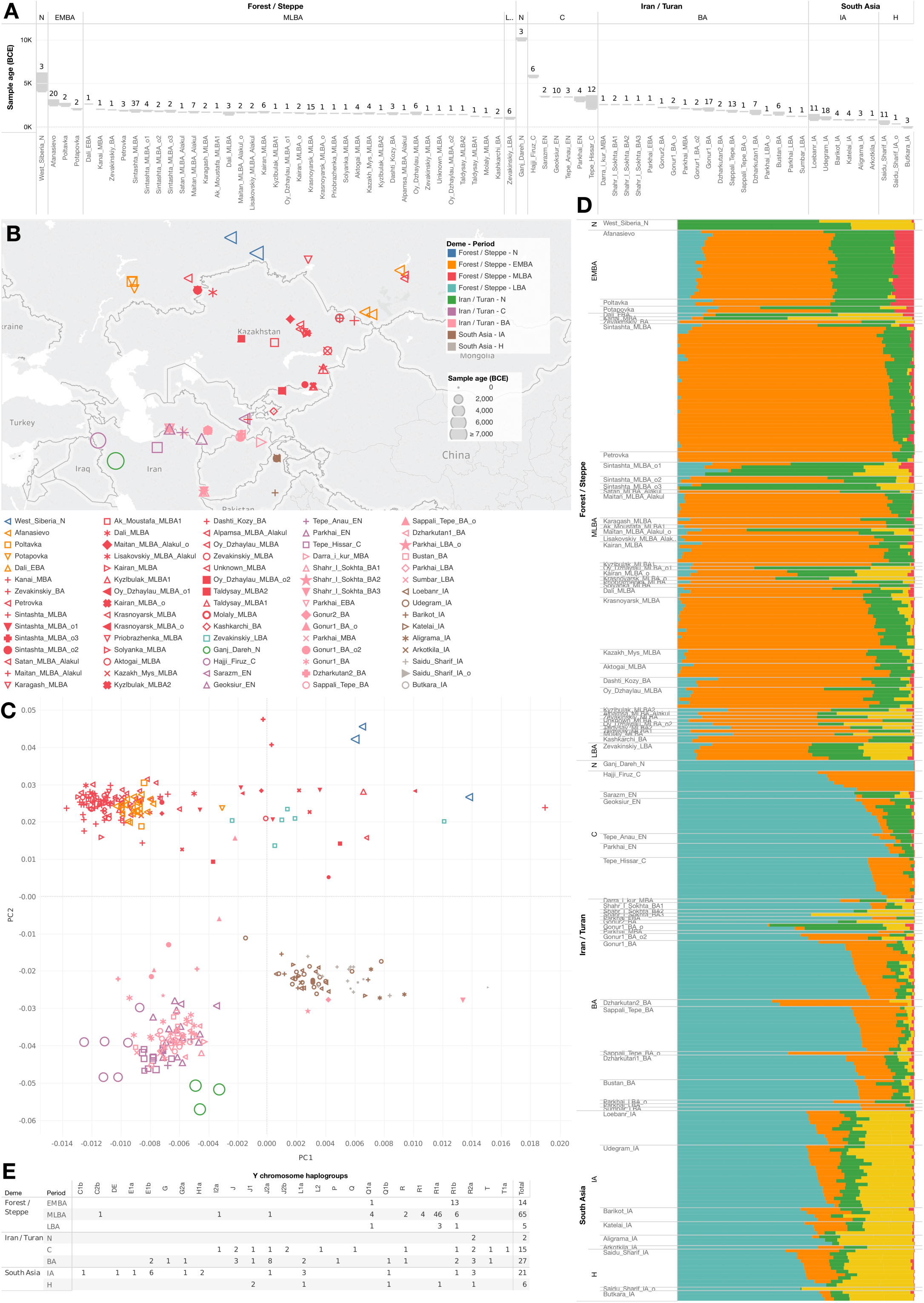
Overview of ancient DNA data. **(A)** Number of newly reported samples passing our analysis thresholds and their date range is shown by site. **(B)** Locations, color-coded by analysis grouping. **(C)** Projections of ancient samples onto PCA axes computed using present-day Eurasians. **(D)** ADMIXTURE analysis, with components maximized in *West_Siberian_HG*, Anatolian agriculturalists, Iranian agriculturalists, indigenous South Asians and *WHG* in blue, orange, teal, red and green, respectively. **(E)** Y-chromosome haplogroups. N, Neolithic; C, Chalcolithic; BA, Bronze Age; IA, Iron Age; H, Historic; E/M/L, Early/Middle/Late; o, outlier.

We carried out principal component analysis (PCA) by projecting the ancient individuals onto the patterns of genetic variation in present-day Eurasians (**Fig. 1**) (*12, 13*). This revealed three major groupings, closely corresponding to the geographic regions of the Forest Zone/Steppe, Iran/Turan and South Asia, a pattern we replicate in ADMIXTURE clustering (*14*). To test formally whether populations differ significantly in their ancestry within regions, we used symmetry-*f*_4_-statistics measuring whether pairs of populations differ in their degree of allele sharing to a third population, and admixture-*f*_3_-statistics to test formally for mixture (**Supplementary Materials**). We tested the fit of mixture models using *qpAdm*, which evaluates whether all possible *f*_4_-statistics relating a set of tested populations to outgroup populations is consistent with mixtures of a pre-specified number of sources and if so estimates proportions of ancestry (*5*). We can model almost every population as a mixture of seven deeply divergent “distal” ancestry sources (usually closely related to populations for which we have data, but in some cases deeply related):

- “Anatolian agriculturalist-related”: represented by 7^th^ millennium BCE western Anatolian agriculturalists (*6*)
- “Western European Hunter-Gatherer (*WHG*)-related”: represented by Mesolithic western Europeans (*5, 10, 15, 16*)
- “Iranian agriculturalist-related”: represented by 8^th^ millennium BCE pastoralists from the Zagros Mountains of Iran (*17, 18*)
- “Eastern European Hunter-Gatherer (*EHG*)-related”: represented by hunter-gatherers from diverse sites in Eastern Europe (*5, 6*)
- “West Siberian Hunter-Gatherer (*West_Siberian_HG)-*related”: a newly documented deep source of Eurasian ancestry represented here by three samples
- “East Asian-related”: represented in this study by Han Chinese
- “Ancient Ancestral South Indian (*AASI*)-related”: a hypothesized South Asian Hunter-Gatherer lineage related deeply to present-day indigenous Andaman Islanders (*19*)

We also used *qpAdm* to identify “proximal” models for each group as mixtures of temporally preceding groups. This often identified multiple alternative models that were equally good fits to the data. These analyses were nevertheless useful because we could identify patterns that were qualitatively consistent across models. The discussion that follows presents an overview of these analyses, while the **Supplementary Materials** presents the full details. **Table 1** summarizes the key findings that emerge from our analysis.

**Table 1.**
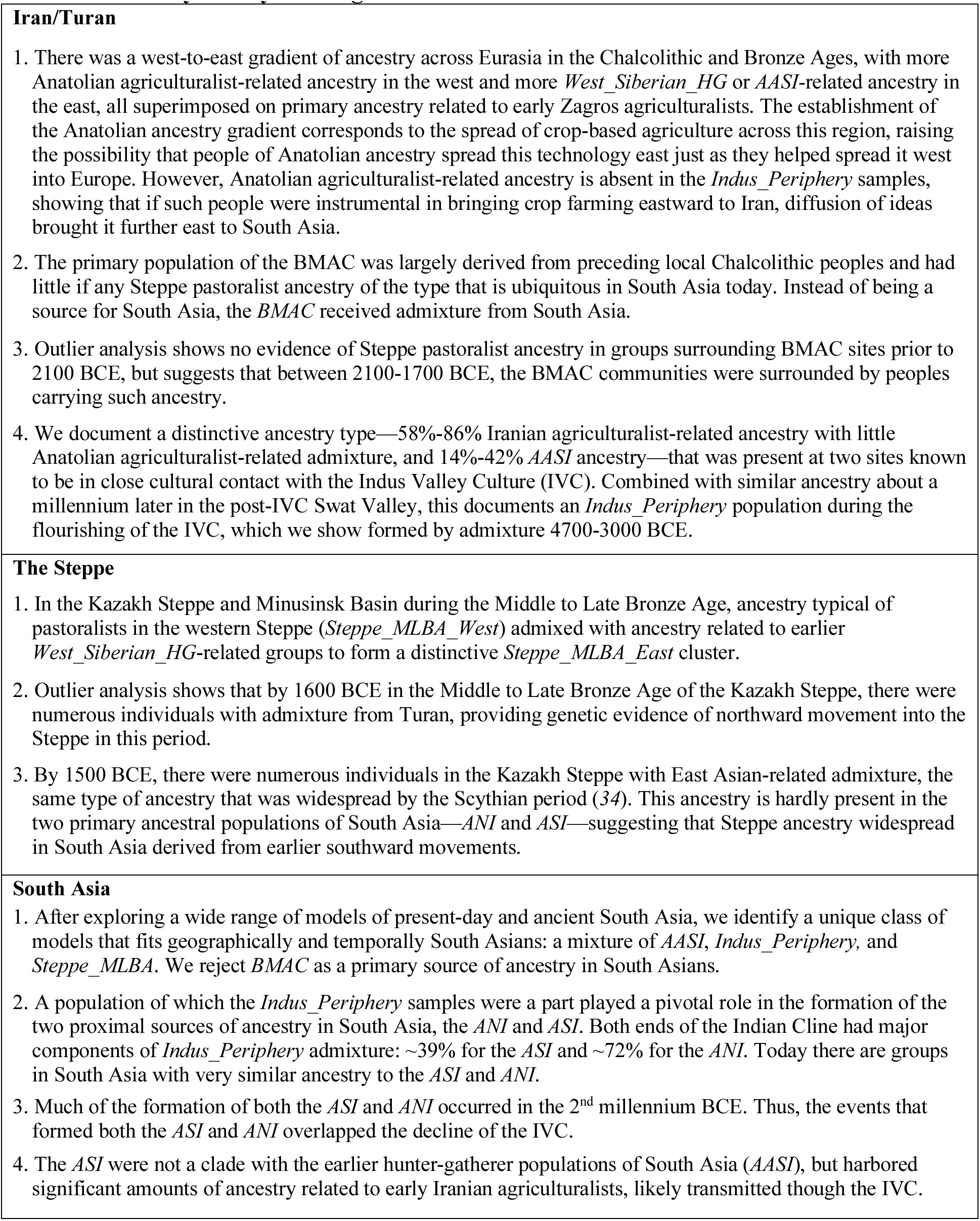
Summary of Key Findings

### Iran/Turan

We analyzed our newly generated data together with previously published data to examine the genetic transformations that accompanied the spread of agriculture eastward from Iran beginning in the 7^th^ millennium BCE (*20, 21*). Our analysis confirms that early Iranian agriculturalists from the Zagros Mountains harbor a distinctive type of West Eurasian ancestry (*17, 18*) (**Fig. 1**), while later groups across a broad geographic region were admixed between this type of ancestry and that related to early Anatolian agriculturalists. (In this paper we use the term “agriculturalists” to refer both to crop cultivation and/or herding, and accordingly refer to the people of the Zagros Mountains who kept domesticated goats as agriculturalists (*17, 22, 23*).) We show that there was a west-to-east cline of decreasing Anatolian agriculturalist-related admixture ranging from ~70% in Chalcolithic Anatolia to ~33% in eastern Iran, to ~3% in far eastern Turan (**Fig. 1; Supplementary Materials**). The timing of the establishment of this cline is consistent with the dates of spread of wheat and barley agriculture from west to east (in the 7^th^ to 6^th^ millennia BCE), suggesting the possibility that individuals of Anatolian ancestry may have contributed to spreading agriculturalist economies not only westward to Europe, but also eastward to Iran (*21, 24, 25*). An increase of Anatolian agriculturalist-related ancestry was also proposed for the Pre-Pottery agriculturalists from the Levant in comparison to the earlier Natufian hunter-gatherers (*17*), further supporting this hypothesis. However, without data on the distribution of Anatolian/Iranian-agriculturalist ancestry in early agriculturalists in Mesopotamia, it is difficult to determine when the cline was established. In the far eastern part of this cline (eastern Iran and Turan) we also detect admixture related to *West_Siberian_HG*, proving that North Eurasian admixture impacted Turan well before the spread of Yamnaya-related Steppe pastoralists (*Steppe_EMBA*).

From Bronze Age Turan, we report 69 ancient individuals (2300-1400 BCE) from four urban sites of the Bactria Margiana Archaeological Complex (BMAC) and its immediate successors. The great majority of individuals fall in a genetic cluster that is similar, albeit not identical, to the preceding groups in Turan in harboring a large proportion of early Iranian agriculturalist-related ancestry (~60% in the *BMAC*) with smaller components of Anatolian agriculturalist-related ancestry (~21%) and *West_Siberian_HG*-related ancestry (~13%) suggesting that the main *BMAC* cluster coalesced from preceding pre-urban populations in Turan (which in turn likely derived from earlier eastward spreads from Iran). The absence in the *BMAC* cluster of the *Steppe_EMBA* ancestry that is ubiquitous in South Asia today—along with *qpAdm* analyses that rule out *BMAC* as a substantial source of ancestry in South Asia (**Fig. 3A**)—suggests that while the BMAC was affected by the same demographic forces that later impacted South Asia (the southward movement of Middle to Late Bronze Age Steppe pastoralists described in the next section), it was also bypassed by members of these groups who hardly mixed with *BMAC* people and instead mixed with peoples further south. In fact, the data suggest that instead of the main *BMAC* population having a demographic impact on South Asia, there was a larger effect of gene flow in the reverse direction, as the main *BMAC* genetic cluster is slightly different from the preceding Turan populations in harboring ~5% of their ancestry from the *AASI*.

**Fig. 3.**
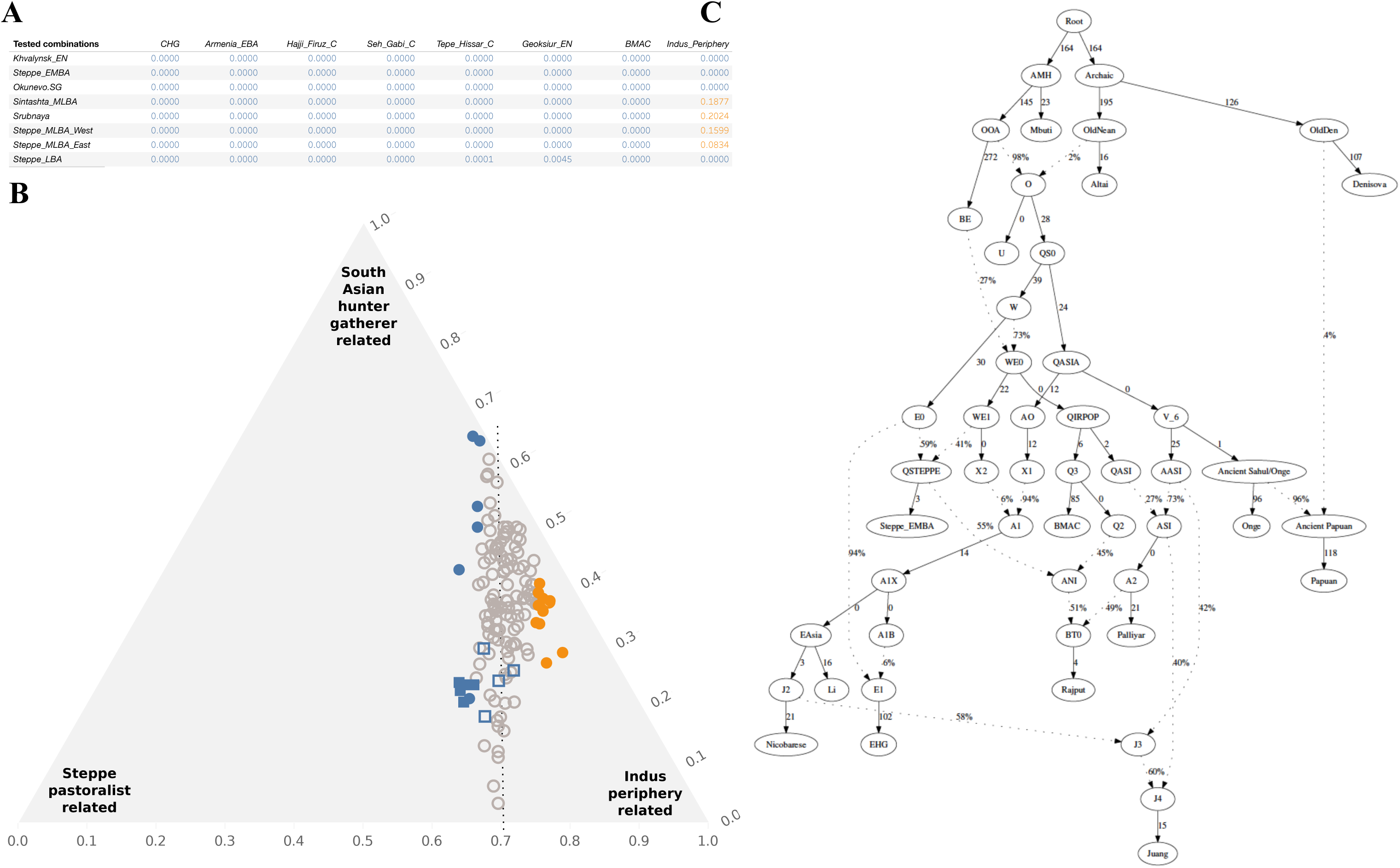
The Genomic Origins of Indians. **(A)** We used *qpAdm* to model four groups that are representative of major sources of South Asian ancestry over the last few thousand years (*Punjabi.DG*, *Mala.DG*, *SPGT*, and *Butkara_IA*) as mixtures of *Onge*, an Iran/Turan-related population, and a Steppe-related group, and report the minimum p-value (highlighting cases at p>0.01). The only working models involve a combination of *Indus_Periphery* and a *Steppe_MLBA* group (note that the *Steppe_MLBA_West* group includes a subset *Sintashta*_*MLBA* and *Srubnaya*). **(B)** For all 140 Indian Cline groups, we give *Maximum A Posteriori* fits for this model. Significant outliers (|Z|>2) are shown, and include a cluster of *Brahmins* (filled circles) and *Bhumihars* (filled squares) with excess Steppe pastoralist-related ancestry compared to others with similar West Eurasian ancestry proportion. **(C)** Admixture graph fit supports Iranian agriculturalist-related admixture into South Asia but no gives evidence of AASI-related admixture into ancient Iran; dotted lines show admixture events.

We also observe outlier individuals at multiple sites, revealing interactions among populations that would be difficult to appreciate without the large sample sizes reported here.

First, around ~2300 BCE in Turan, we observe two outliers at the BMAC site of Gonur with *West_Siberian_HG*-related ancestry of a type that we observe at multiple sites in Kazakhstan over the preceding and succeeding millennia. The most plausible explanation is that this ancestry is that of indigenous populations associated with the Kelteminar culture, the native hunter-gatherers of the region who covered a vast area of Central Asia before the BMAC (*26*). Future ancient DNA data from Kelteminar contexts will make it possible to determine whether it is indeed the case that the genetic ancestry of Kelteminar people was similar to that of *West_Siberia_HG.* Importantly, in the 3^rd^ millenium BCE we do not find any individuals with ancestry derived from Yamnaya-related Steppe pastoralists in Turan. Thus, *Steppe_EMBA* ancestry was not yet widespread across the region.

Second, between 2100-1700 BCE, we observe BMAC outliers from three sites with *Steppe_EMBA* ancestry in the admixed form typically carried by the later Middle to Late Bronze Age Steppe groups (*Steppe_MLBA*). This documents a southward movement of Steppe ancestry through this region that only began to have a major impact around the turn of the 2^nd^ millennium BCE.

Third, between 3100-2200 BCE we observe an outlier at the BMAC site of Gonur, as well as two outliers from the eastern Iranian site of Shahr-i-Sokhta, all with an ancestry profile similar to 41 ancient individuals from northern Pakistan who lived approximately a millennium later in the isolated Swat region of the northern Indus Valley (1200-800 BCE). These individuals had between 14-42% of their ancestry related to the AASI and the rest related to early Iranian agriculturalists and *West_Siberian_HG.* Like contemporary and earlier samples from Iran/Turan we find no evidence of Steppe-pastoralist-related ancestry in these samples. In contrast to all other Iran/Turan samples, we find that these individuals also had negligible Anatolian agriculturalist-related admixture, suggesting that they might be migrants from a population further east along the cline of decreasing Anatolian agriculturalist ancestry. While we do not have access to any DNA directly sampled from the Indus Valley Civilization (IVC), based on (a) archaeological evidence of material culture exchange between the IVC and both BMAC to its north and Shahr-i-Sokhta to its east (*27*), (b) the similarity of these outlier individuals to post-IVC Swat Valley individuals described in the next section (*27*), (c) the presence of substantial *AASI* admixture in these samples suggesting that they are migrants from South Asia, and (d) the fact that these individuals fit as ancestral populations for present-day Indian groups in *qpAdm* modeling, we hypothesize that these outliers were recent migrants from the IVC. Without ancient DNA from individuals buried in IVC cultural contexts, we cannot rule out the possibility that the group represented by these outlier individuals, which we call *Indus_Periphery*, was limited to the northern fringe and not representative of the ancestry of the entire Indus Valley Civilization population. In fact, it was certainly the case that the peoples of the Indus Valley were genetically heterogeneous as we observe one of the *Indus_Periphery* individuals having ~42% *AASI* ancestry and the other two individuals having ~14-18% *AASI* ancestry (but always mixes of the same two proximal sources of *AASI* and Iranian agriculturalist-related ancestry). Nevertheless, these results show that *Indus_Periphery* were part of an important ancestry cline in the wider Indus region in the 3^rd^ millennium and early 2^nd^ millennium BCE. As we show in what follows, peoples related to this group had a pivotal role in the formation of subsequent populations in South Asia.

Using a newly developed approach for estimating dates of admixture in ancient genomes (an adaptation of a previous method to measure ancestry covariance among pairs of neighboring positions in the genome; **Supplementary Materials**), we estimate that the time of admixture between Iranian agriculturalist-related ancestry and AASI ancestry in the three *Indus_Periphery* samples was 53 ± 15 generations ago on average, corresponding to a 95% confidence interval of about 4700-3000 BCE assuming 28 years per generation (*28*). This places a minimum date on the first contact between these two types of ancestries.

### The Steppe

Three individuals from the West Siberian forest zone with direct dates ranging from 6200 BCE to 4000 BCE play an important role in this study as they are representatives of a never-before-reported mixture of ancestry that we call *West_Siberian_HG*: ~30% derived from *EHG*, ~50% from Ancestral North Eurasians (defined as being related deeply to 22000-15000 BCE Siberians (*29, 30*)), and ~20% related to present-day East Asians. This ancestry type also existed in the southern Steppe and in Turan, as it formed about 80% of the ancestry of an early 3^rd^ millennium BCE agro-pastoralist from Dali, Kazakhstan, and also contributed to multiple outlier individuals from 2^nd^ millennium sites in Kazakhstan and Turan (**Fig. 2**).

**Fig. 2.**
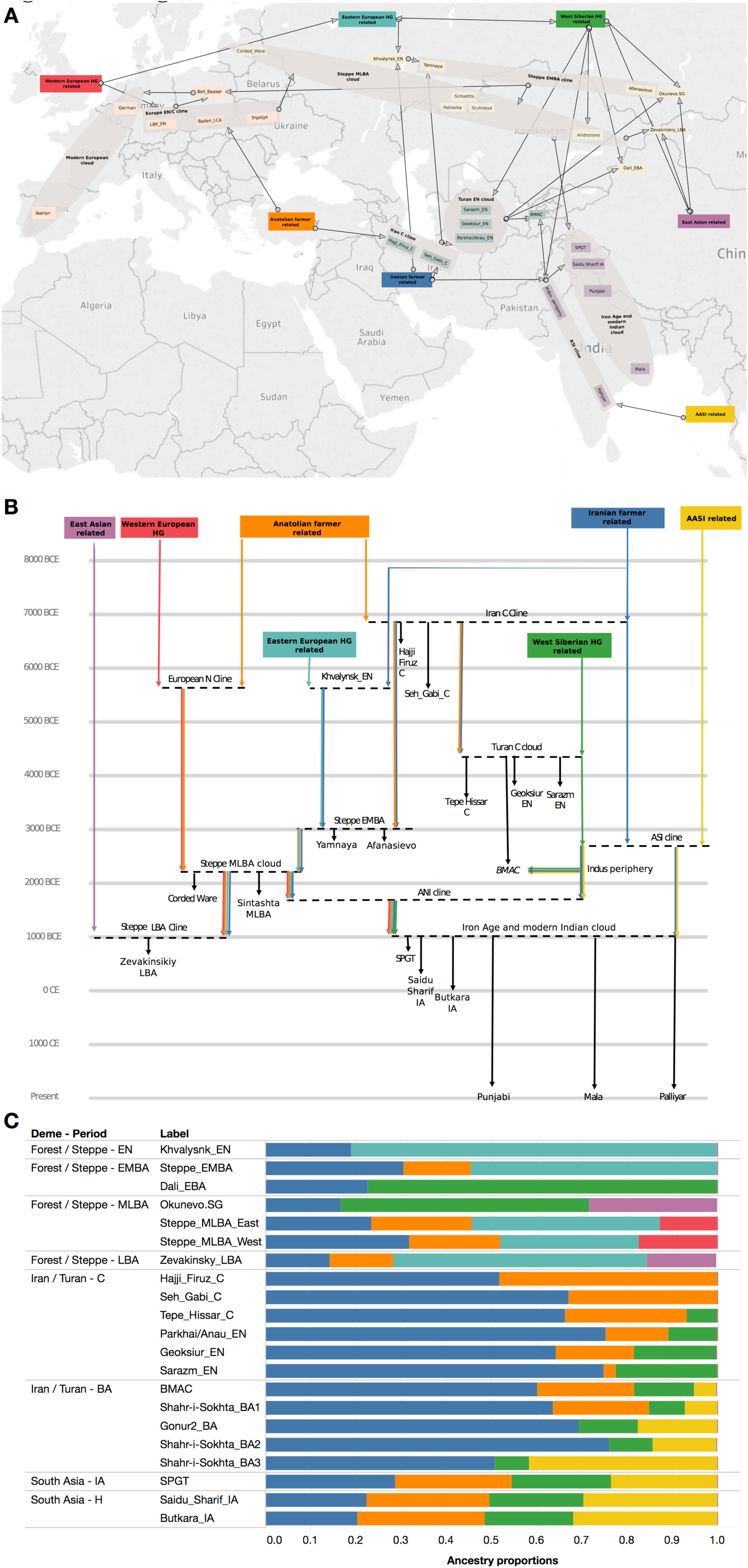
Modeling results. **(A)** Admixture events originating from 7 “Distal” populations leading to the formation of the modern Indian cloud shown geographically. Clines or 2-way mixtures of ancestry are shown in rectangles, and clouds (3-way mixtures) are shown in ellipses. **(B)** A schematic model of events originating from 7 “Distal” populations leading to the formation of the modern Indian cline, shown chronologically. **(C)** Admixture proportions as estimated using *qpAdm* for populations reflected in A and B.

Using the *West_Siberian_HG* individuals as a reference population along with other pre-Chalcolithic groups that have been previously reported in the ancient DNA literature, we document the presence of a genetically relatively homogeneous population spread across a vast region of the eastern European and trans-Ural Steppe between 2000-1400 BCE (*Steppe_MLBA*) (*17*). Many of the samples from this group are individuals buried in association with artifacts of the Corded Ware, Srubnaya, Petrovka, Sintashta and Andronovo complexes, all of which harbored a mixture of *Steppe_EMBA* ancestry and ancestry from European Middle Neolithic agriculturalists (*Europe_MN*). This is consistent with previous findings showing that following westward movement of eastern European populations and mixture with local European agriculturalists, there was an eastward reflux back beyond the Urals (*6, 16, 31*). Our new dataset enhances our understanding of the *Steppe_MLBA* cluster by including many sites in present-day Kazakhstan and as far east as the Minusinsk Basin of Russia—and in doing so allows us to appreciate previously undetected substructure. All previously reported samples fall into a subcluster we call *Steppe_MLBA_West* that harbors ~26% *Europe_MN* ancestry and ~74% *Steppe_EMBA* ancestry. With our newly reported data we now also detect a previously unappreciated subcluster, *Steppe_MLBA_East,* which is significantly differentiated (p=7×10^−6^ from *qpAdm*), with ~8% *West_Siberian_HG*-related ancestry and proportionally less of the other ancestry components, suggesting that people carrying *Steppe_MLBA_West* ancestry admixed with *West_Siberian_HG*-related peoples as they spread further east.

As in Iran/Turan, the outlier individuals provide key additional information.

First, our analysis of 50 newly reported individuals from the Kamennyi Ambar V cemetery from the Sintashta culture reveals three groups of outliers, in addition to the main cluster of 40 individuals. These outliers have elevated proportions of *Steppe_EMBA*, *West_Siberian_HG* or *East Asian-*related ancestry (and direct dates that are contemporaneous with the other individuals), thereby showing that this fortified site harbored people of diverse ancestries living side-by-side.

Second, samples from three sites from the southern and eastern end of the Steppe dated to 1600-1500 BCE (Dashti-kozy, Taldysay and Kyzlbulak) show evidence of significant admixture from Iranian agriculturalist-related populations, demonstrating northward gene flow from Turan into the Steppe at the same time as there was southward movement of *Steppe_MLBA* ancestry through Turan and into South Asia. These findings are consistent with evidence of a high degree of human mobility both to the north and south along the Inner Asian Mountain Corridor (*32, 33*).

Third, we observe samples from multiple sites dated to 1700-1500 BCE (Maitan, Kairan, Oy_Dzhaylau and Zevakinsikiy) that derive up to ~25% of their ancestry from a source related to present-day East Asians and the remainder from *Steppe_MLBA*. A similar ancestry profile became widespread in the region by the Late Bronze Age, as documented by our time transect from Zevakinsikiy and samples from many sites dating to 1500-1000 BCE, and was ubiquitous by the Scytho-Sarmatian period in the Iron Age (*34*). This observation decreases the probability that populations in the 1^st^ millennium BCE and 1^st^ millennium CE—including Scythians, Kushans, and Huns, sometimes suggested as sources for the Steppe ancestry influences in South Asia today (*17*)—contributed to the majority of South Asians, which have negligible East Asian ancestry in our analysis. It is possible that there were unsampled groups in Central Asia with negligible East Asian admixture that could have migrated later to South Asia. However, at least some (possibly all) of the Steppe pastoralist ancestry in South Asia owes its origins to southward pulses in the 2^nd^ millennium BCE, as indeed we prove directly through our observation of this ancestry in the Swat Iron Age individuals dating to ~1000 BCE (discussed further below).

### South Asia

Previous work has shown that the Indian Cline—a gradient of different proportions of West Eurasian related ancestry in South Asia—can be well modeled as having arisen from a mixture of two statistically reconstructed ancestral populations (the *ANI* and the *ASI*), which mixed mostly after 2000 BCE (*35, 36*). Ancient DNA analysis has furthermore revealed that the populations along the Indian Cline actually descend more deeply in time from at least three ancestral populations (*17*), with ancestry from groups related to early Iranian agriculturalists, *Steppe_EMBA*, and *Onge*.

To shed light on the mixture events that transformed this minimum of three ancestral populations into two (the *ANI* and *ASI*), we used *qpAdm* to search for triples of source populations—the *AASI*, all sampled ancient Iran/Turan-related groups, and all sampled ancient Steppe groups— that could fit as sources for South Asians. As South Asian test populations we used an Indian Cline group with high ANI ancestry (*Punjabi.DG*), one with high ASI ancestry (*Mala.DG*), early Iron Age Swat Valley samples *(Swat Protohistoric Grave Type - SPGT),* and Early Historic Swat Valley samples *(Butkara_IA*). **Fig. 3A** shows that the only models that fit all four test South Asians groups are combinations that involve the *AASI*, *Indus_Periphery* and *Steppe_MLBA* (in the analyses that follow, we therefore pooled the *Steppe_MLBA*). The evidence that the *Steppe_MLBA* cluster is a plausible source for the Steppe ancestry in South Asia is also supported by Y chromosome evidence, as haplogroup R1a which is of the Z93 subtype common in South Asia today (*37, 38*) was of high frequency in *Steppe_MLBA* (68%) (*16*), but rare in *Steppe_EMBA* (absent in our data).

To obtain a richer understanding of the ancestry of the entire Indian Cline, we took advantage of previously published genome-wide data from 246 ethnographically diverse groups from South Asia (*11*), from which we sub-selected 140 groups that fall on a clear gradient in PCA to represent the Indian Cline (the other groups either fall off the cline due to additional African or East Asian-related ancestry or had small sample size or heterogeneous ancestry). The per-group *qpAdm* estimates for the proportions of ancestry from these three sources are statistically noisy. We therefore developed new methodology that allows us to jointly fit the data from all Indian Cline groups within a hierarchical model. The analysis confirms that the great majority of all groups on the Indian Cline can be jointly modeled as a mixture of two populations, and the analysis also produces an estimate of the functional relationship between the ancestry components. Setting *Steppe_MLBA* to its smallest possible proportion of zero to estimate the minimum fraction of *Indus_Periphery* ancestry that could have existed in the *ASI,* we obtain ~39%. Setting *AASI* to its smallest possible proportion of zero to estimate the maximal fraction of *Indus_Periphery* ancestry that could have existed in the *ANI,* we obtain ~72%. In fact, we find four tribal groups from southern India (*Palliyar, Ulladan, Malayan,* and *Adiyan*) with close to the maximum mathematically allowed proportion of *Indus_Periphery-*related ancestry, and we find a population in northern Pakistan (*Kalash*) with close to the minimum. Thus, nearly unmixed descendants of the *ASI* and *ANI* exist as isolated groups in South Asia today.

We built an admixture graph using *qpGraph* co-modeling *Palliyar* (as a representative of the *ASI*) and *Juang* (an Austroasiatic speaking group in India with low West Eurasian-relatedness), and show that it fits when the ASI have ~27% Iranian agriculturalist-related ancestry and the *Juang* also harbor ancestry from an AASI population without Iranian admixture (**Fig. 3**). This model is also notable in showing that early Iranian agriculturalists fit without *AASI* admixture, and thus the patterns we observe are driven by gene flow into South Asia and not the reverse (**Fig. 3; Supplementary Materials**). The fitted admixture graph also reveals that the deep ancestry of the indigenous hunter-gather population of India represents an anciently divergent branch of Asian human variation that split off around the same time that East Asian, *Onge* and *Australian* aboriginal ancestors separated from each other. This finding is consistent with a model in which essentially all the ancestry of present-day eastern and southern Asians (prior to West Eurasian-related admixture) derives from a single eastward spread, which gave rise in a short span of time to the lineages leading to *AASI*, East Asians, *Onge*, and *Australians* (*19*).

Using admixture linkage disequilibrium, we estimate a date of 107 ± 11 generations ago for the Iranian agriculturalist and AASI-related admixture in the *Palliyar*, corresponding to a 95% confidence interval of 1700-400 BCE assuming 28 years per generation (*28*). This date is consistent with a previous estimate of 110 ± 12 generations ago for the *Kalash* (*39*). These results suggest that the *ASI* and *ANI* were both largely unformed at the beginning of the 2^nd^ millennium BCE, and imply that the *ASI* may have formed in the course of the spread of West Asian domesticates into peninsular India beginning around 3000 BCE (where they were combined with local domesticates to form the basis of the early agriculturalist economy of South India (*40*)), or alternatively in association with eastward spread of material culture from the Indus Valley after the IVC declined (*41*). Further evidence for a Bronze Age formation of the *ASI* comes from our analysis of Austroasiatic-speaking groups in India such as *Juang,* who have a higher ratio of *AASI*-to-Iranian agriculturalist-related ancestry than the *ASI* (**Fig. 3, Supplementary Materials**). Austroasiatic speakers likely descend from populations that arrived in South Asia in the 3^rd^ millennium BCE (based on hill cultivation systems associated with the spread of Austroasiatic languages (*20*)), and our genetic results show that when Austroasiatic speakers arrived they mixed with groups with elevated ratios of *AASI*-to Iranian-agriculturalist-related ancestry than are found in the *ASI*, showing that the *ASI* had not yet overspread peninsular India.

Finally, we examined our Swat Valley time transect from 1200 BCE to 1 CE. While the earliest group of samples (*SPGT*) is genetically very similar to the *Indus_Periphery* samples from the sites of *Gonur* and *Shahr-i-Sokhta,* they also differ significantly in harboring *Steppe_MLBA* ancestry (~22%). This provides direct evidence for *Steppe_MLBA* ancestry being integrated into South Asian groups in the 2^nd^ millennium BCE, and is also consistent with the evidence of southward expansions of *Steppe_MLBA* groups through Turan at this time via outliers from the main *BMAC* cluster from 2000-1500 BCE. Later samples from the Swat time transect from the 1^st^ millennium BCE had higher proportions of Steppe and AASI derived ancestry more similar to that found on the Indian Cline, showing that there was an increasing percolation of Steppe derived ancestry into the region and additional admixture with the ASI through time.

### Implications for Archaeology and Linguistics

Our evidence that a population with both Iranian agriculturalist and South Asian hunter gatherer ancestry (*Indus_Periphery*) was established in the 3^rd^ millennium BCE—and that its Iranian agriculturalist-related and *AASI* ancestry sources mixed at an average time of around 4700-3000 BCE—shows that this type of Iranian agriculturalist-related ancestry must have reached the Indus Valley by the 4^th^ millennium BCE. However, it is very possible that Iranian agriculturalist-related ancestry was widespread in South Asia even earlier, as wheat and barley agriculture as well as goat and sheep herding spread into South Asia after the 7^th^ millennium BCE, as attested at sites such as Mehrgarh in the hills surrounding the Indus Valley (*20, 21*), and these domesticates could have been carried by movements of people. Regardless of when these agricultural species arrived, the genetic data show that *Indus_Periphery-*related ancestry contributed in large proportions to both the *ANI* and *ASI*, and that these two groups both formed in the 2^nd^ millennium BCE, overlapping the decline of the IVC and major changes in settlement patterns in the northern part of the Indian subcontinent (*41*). A parsimonious hypothesis is that as *Steppe_MLBA* groups moved south and mixed with *Indus_Periphery*-related groups at the end of the IVC to form the *ANI,* other *Indus_Periphery*-related groups moved further south and east to mix with *AASI* groups in peninsular India to form the *ASI*. This is consistent with suggestions that the spread of the IVC was responsible for dispersing Dravidian languages (*42–44*), although scenarios in which Dravidian languages derive from pre-Indus languages of peninsular India are also entirely plausible as *ASI* ancestry is mostly derived from the *AASI*.

Our results also shed light on the question of the origins of the subset of Indo-European languages spoken in India and Europe (*45*). It is striking that the great majority of Indo-European speakers today living in both Europe and South Asia harbor large fractions of ancestry related to Yamnaya Steppe pastoralists (corresponding genetically to the *Steppe_EMBA* cluster), suggesting that “Late Proto-Indo-European”—the language ancestral to all modern Indo-European languages—was the language of the Yamnaya (*46*). While ancient DNA studies have documented westward movements of peoples from the Steppe that plausibly spread this ancestry to Europe (*5, 31*), there has not been ancient DNA evidence of the chain of transmission to South Asia. Our documentation of a large-scale genetic pressure from *Steppe_MLBA* groups in the 2^nd^ millennium BCE provides a prime candidate, a finding that is consistent with archaeological evidence of connections between material culture in the Kazakh middle-to-late Bronze Age Steppe and early Vedic culture in India (*46*).

Our analysis also provides an entirely new line of evidence for a linkage between Steppe ancestry and Indo-European culture. When we used *qpAdm* to test if a mixture of ANI and ASI is a fit to the data for all 140 Indian Cline groups, we found 10 groups with poor fits and a significantly elevated ratio of *Steppe_MLBA-*to *Indus_Periphery-*related ancestry compared to the expectation for the model (Z≥3). We found the strongest two signals in *Brahmin_Tiwari* (p=2×10^−5^) and *Brahmin_UP* (p=4×10^−5^), and more generally there was a striking enrichment of a Z≥3 signals in groups of traditionally priestly status in northern India (57% of groups with Z≥3 were *Brahmins* or *Bhumihars* even though these groups comprised only 11% of the 74 groups we analyzed in northern India). Although the enrichment for Steppe ancestry is not found in the southern Indian groups, the Steppe enrichment in the northern groups is striking as *Brahmins* and *Bhumihars* are among the traditional custodians of texts written in early Sanskrit. A possible explanation is that the influx of *Steppe_MLBA* ancestry into South Asia in the mid-2^nd^ millennium BCE created a meta-population of groups with different proportions of Steppe ancestry, with ones having relatively more Steppe ancestry having a central role in spreading early Vedic culture. Due to strong endogamy in South Asia—which has kept some groups isolated from their neighbors for thousands of years (*35*)—some of this substructure within Indian population still persists.

We finally highlight a remarkable parallel between the prehistory of two sub-continents of Eurasia: South Asia and Europe. In both regions, West Asian agricultural technology spread from an origin in the Near East in the 7^th^ and 6^th^ millennia BCE (**Fig. 4**). In South Asia this occurred via the Iranian plateau, and in Europe via western Anatolia, with the technological spreads mediated in both cases by movements of people. An admixed population was then formed by the mixing of incoming agriculturalists and resident hunter-gatherers—in South Asia eventually giving rise to the *Indus_Periphery* and *ASI* and in Europe the Middle Neolithic genetic cluster *Europe_MN*. In both Europe and South Asia, populations related to the *Yamnaya* Steppe pastoralists arrived after this agriculturalist and hunter-gatherer admixture took place, interacting with local populations to produce mixed groups, which then mixed further with already resident agriculturalist populations to produce genetic groupings such as those found associated with Corded Ware and central European Bell Beaker artifacts in much of Europe, and the *ANI* genetic cluster in South Asia. These mixed groups then mixed further to produce the major gradients of ancestry in both regions. Future studies of populations from South Asia and the linguistically related Iranian world will extend and add nuance to the model presented here.

**Fig. 4.**
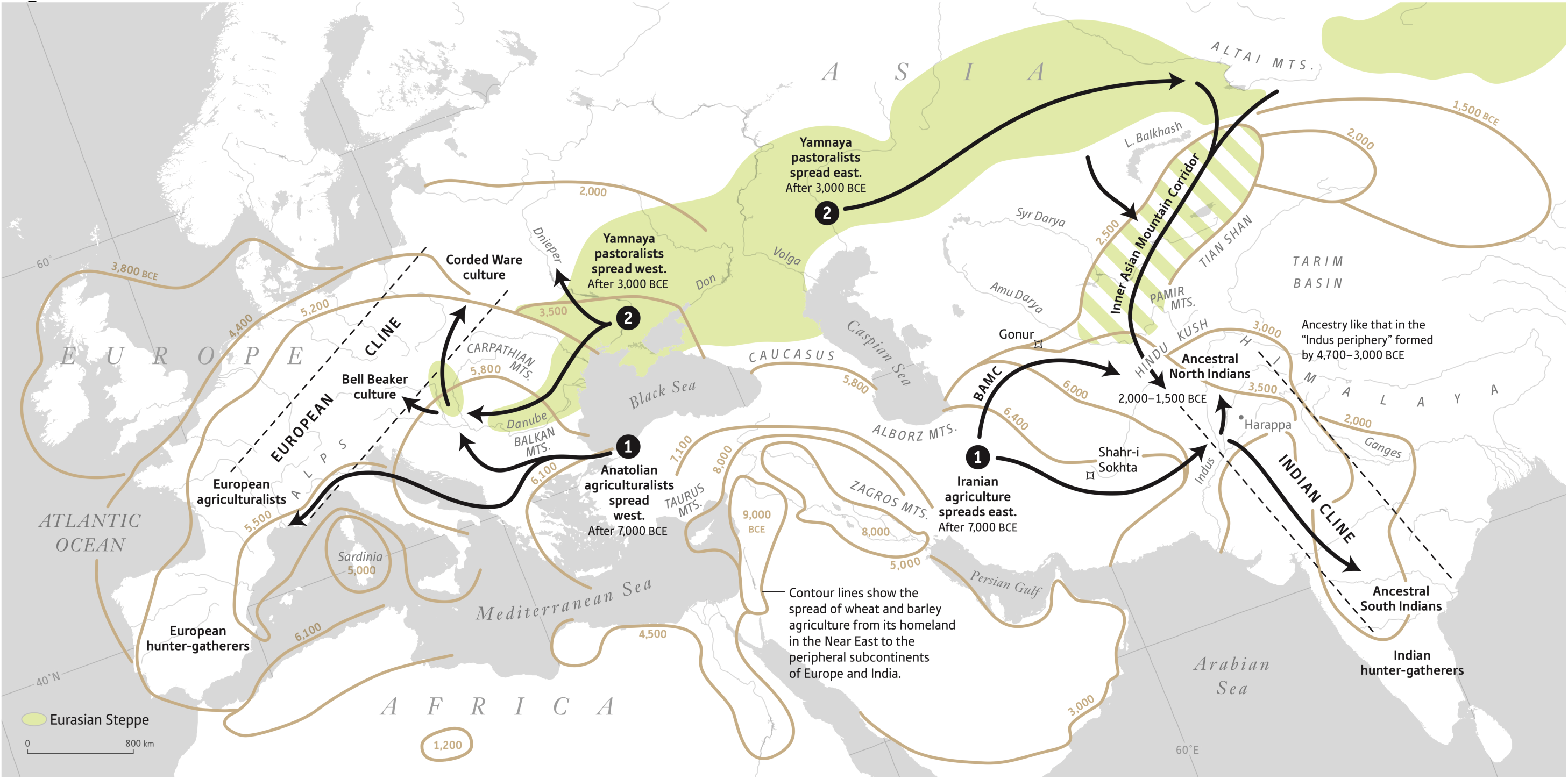
A Tale of Two Subcontinents. The prehistory of South Asia and Europe are parallel in both being impacted by two successive spreads, the first from the Near East after 7000 BCE bringing agriculturalists who mixed with local hunter-gatherers, and the second from the Steppe after 3000 BCE bringing people who spoke Indo-European languages and who mixed with those they encountered during their migratory movement. Mixtures of these mixed populations then produced the rough clines of ancestry present in both South Asia and in Europe today (albeit with more variable proportions of local hunter-gatherer-related ancestry in Europe than in India), which are (imperfectly) correlated to geography. The plot shows in contour lines the time of the expansion of Near Eastern agriculture. Human movements and mixtures, which also plausibly contributed to the spread of languages, are shown with arrows.

## Materials and Methods

### Ancient DNA Laboratory Work

We prepared powder from human skeletal remains either at field sites using a method for extracting petrous bone powder by drilling directly from the cranial base (*47*), or in dedicated clean rooms at Harvard Medical School, University College Dublin, or the Max Planck Institute for Evolutionary Anthropology in Leipzig Germany.

All the molecular work except for that of a single sample (Darra-i-Kur) was carried out at Harvard Medical School (HMS). At HMS, we extracted DNA using a method that is optimized to retain small DNA fragments (*1, 2*). We converted the DNA into a form that could be sequenced using a double-stranded library preparation protocol, usually pre-treating with the enzyme Uracil-DNA Glycosylase (UDG) to reduce the characteristic cytosine-to-thymine errors in ancient DNA (*4*). For some libraries, we substituted the MinElute columns used for cleaning up reactions with magnetic beads, and the MinElute column-based PCR cleanup at the end of library preparation with SPRI beads (*48, 49*). We enriched the libraries both for sequences overlapping mitochondrial DNA (*50*), and for sequences overlapping about 1.24 million nuclear targets after two rounds of enrichment (*5, 6, 8*). We sequenced the enriched products on an Illumina NextSeq500 using v.2 150 cycle kits for 2×76 cycles and 2×7 cycles, and sequenced up to the point that the expected number of new SNPs covered per 100 additional read pairs sequenced was approximately less than 1.

To analyze the data computationally, we separated read pairs into individuals based on searching for the expected two indices and two barcodes, allowing up to one mismatch from the expected sequence in each case. We removed adapters and merged together sequences requiring a 15 base pair overlap (allowing up to one mismatch), using a modified version of Seqprep https://github.com/jstjohn/SeqPrep), which takes the highest quality base in the merged regions. We mapped the resulting single-ended sequences were mapped to the GRCh37 human reference (from the 1000 Genomes project) using the *samse* command of the Burrows-Wheeler Aligner tool (*BWA*) (version 0.6.1) (*51*). We trimmed two nucleotides from the end of each sequence, and then randomly selected a single sequence at each site covered by at least one sequence in each individual to represent their genotype at that position (“pseudo-haploid” genotyping). For each sample we generated “pseudo-haploid” calls at the 1.24 million target sites, selecting sequences that have a minimum mapping quality of MAPQ≥10, restricting to nucleotides with a minimum base quality of 20, and trimming 2 base pairs from each end of the reads.

For Darra-i-Kur, we prepared a single-stranded DNA library (L5082) at the Max-Planck-Institute for Evolutionary Anthropology (MPI-EVA) in Leipzig, Germany, as part of a previous project (*52*). The previous study only analyzed mitochondrial DNA, and for the current study, the library was enriched for molecules overlapping target the same panel of 1.24 million nuclear targets using two rounds of hybridization capture (*5, 6, 8*). We sequenced the enriched libraries on 2 lanes of an Illumina HiSeq2500 platform in a double index configuration (2×76 cycles) (*53*), and we called sites using *FreeIbis* (*54*). We merged overlapping paired-end and trimmed using *leeHom* (*55*). We used *BWA* to align the captured data to the human reference genome (GRCh37 from the 1000 Genomes project) (*51*). Only sequences showing a perfect match to the expected index combination were retained for downstream analyses.

We assessed evidence for ancient DNA authenticity by measuring the rate of damage in the first nucleotide (flagging individuals as potentially contaminated if they had a less than 3% cytosine to thymine substitution rate in the first nucleotide for a UDG-treated library and less than 10% substitution rate for a non-UDG-treated library). We used *contamix* to determine evidence of contamination based on polymorphism in mitochondrial DNA (*56*), and ANGSD to determine evidence of contamination based on polymorphism on the X chromosome in males (*57*).

### Principal component analysis (PCA)

We carried out PCA using the *smartpca* package of *EIGENSOFT* 7.2.1 (*13*). We used default parameters and added two options (lsqproject:YES and numoutlieriter:0 options) in order to project our ancient samples onto the PCA space. We used two basis sets for the projection: the first based on 1,340 present-day Eurasians genotyped on the Affymetrix Human Origins array, and the second based on a subset of 991 present-day West Eurasians (*5, 10, 58*). These projections are shown repeatedly in the **Supplementary Materials**, and the whole-Eurasian projection is shown in **Fig. 1**. As part of this analysis, we also computed the F_ST_ between groups using the parameters inbreed:YES and fstonly:YES.

### ADMIXTURE clustering analysis

Using PLINK2 (*59*), we first pruned our dataset using the -- geno 0.7 option to ensure that we only performed our analysis on sites that had at least 70% of samples with a called genotype. We then ran ADMIXTURE (*14*) with 10 replicates, reporting the replicate with the highest likelihood. We show results for K=6 in **Fig. 1**, as we found in practice that this provides the most resolution for disambiguating the sources of pre-Chalcolithic ancestry in our newly reported samples.

### f-statistics

We used the *qp3pop* and *qpDstat* packages in ADMIXTOOLS to compute *f*_3_-statistics and *f*_4_-statistics. We used the inbreed:YES parameter to compute *f*_3_-statistics as a test for admixture with an ancient population as a target, with all published and newly reported ancient genomes as sources. Using the f4Mode:YES parameter in *qpDstat,* we also computed two sets of *f*_4_-symmetry statistics to evaluate if pairs of populations are consistent with forming a clade relative to a comparison population. The first is a statistic where we compare all possible pairs of newly reported ancient groups (*Reported1* and *Reported2*) to a panel of *Test* populations that encompass diverse pre-Chalcolithic and more widespread genetic variation (*Test* is one of *Iran_Ganj_Dareh_Neolithic, Karelia_HG, Han, Onge, LBK_EN, AfontovaGora3, Ukraine_Mesolithic*). Thus, we compute a statistic of the form *f*_4_(*Reported1, Reported2; Test, Mbuti* African outgroup). The second is a comparison of each newly reported group in turn against all possible pairs of *Test* populations, using statistics of the form *f*_4_(*Test1*, *Test2*; *Reported*, *Mbuti*).

### Formally modeling admixture history

We used the *qpAdm* methodology (*5*) in the ADMIXTOOLS package to estimate the proportions of ancestry in a *Test* population deriving from a mixture of *N* ‘reference’ populations by exploiting (but not explicitly modeling) shared genetic drift with a set of ‘Outgroup’ populations. We set the details:YES parameter, which reports a normally distributed Z-score for the fit (estimated with a block jackknife).

### Hierarchical model of the Indian Cline

We used *qpAdm* as described above to obtain estimates for the proportion of Steppe-related, Iranian agriculturalist-related and *AASI*-related ancestries and their relevant covariance matrices for each population on the Indian cline. We then jointly modeled hese estimates using a bivariate normal model (since the three proportions sum to 100%) and nferred the mean and covariance of the two parameters across all samples on the Indian cline using maximum likelihood estimation. Then, using this inferred matrix, we tested whether the cline could be modeled by a mixture of two populations, the *ANI* and the *ASI*, in two ways. First, we examined whether the covariance matrix is singular, implying that knowledge of one estimated proportion of ancestry of one of the ancestry components revealed knowledge of the other two, as expected in a two-way mixture. Second, if we were able to establish that this was the case, we examined the difference between the expected and observed ratios of the ancestry proportions of individual populations in this generative model obtained from fitting all the populations simultaneously. This process resulted in a handful of populations deviating from expectation, as discussed in the main text and **Supplementary Materials**.

## Supplementary Materials

Materials and Methods

Online Tableau Server for visualizing data.

Data S1-S3

## Acknowledgments

We are grateful to Richard Meadow and Ajita Patel for critical comments. We are grateful to the Minusinsk Regional Museum of N. M. Martyanov for sharing some of the skeletal samples analyzed in this study. We are grateful to Orazak Ismagulov and Ainagul Ismagulova for facilitating access to some of the Kazakh material.

## Data and materials availability

All sequencing data are available from the European Nucleotide Archive, accession number XXXXXXXX [to be made available on publication]. Genotype data obtained by random sampling of sequences at approximately 1.24 million analyzed positions or at approximately 600,000 positions (when merged with genotyping data from diverse present-day individuals) are available to researchers who write David Reich a signed letter containing the following text: “For the data that is indicated as “signed letter only” (a) I will not distribute the data outside my collaboration; (b) I will not post the data publicly; (c) I will make no attempt to connect the genetic data to personal identifiers for the samples; (d) I will use the data only for studies of population history; (e) I will not use the data for any selection studies; (f) I will not use the data for medical or disease-related analyses; (g) I will not use the data for commercial purposes.’’

